# Data-driven representations using deep network-coherent methylation autoencoders to identify robust disease and risk factor signatures

**DOI:** 10.1101/2023.03.07.531501

**Authors:** David Martínez-Enguita, Sanjiv K. Dwivedi, Rebecka Jörnsten, Mika Gustafsson

## Abstract

Data analysis in systems medicine often employs knowledge-driven approaches, which leverage prior biological insights to guide and inform the study of large omic sets. However, the current state of knowledge in biology is still partial and biased. For example, cancer-associated genes are overrepresented in protein interaction networks. As a result, these approaches may fail to capture novel or unexpected phenomena. In this study, we present a data-driven workflow for the functional analysis of large DNA methylation data using deep autoencoders with biologically relevant latent embeddings (network-coherent autoencoders, NCAEs). We observed an increasing gene co-localization gradient, consistent with the human interactome, within the learned representation of a deep methylation autoencoder. We showcased the capacity of this coherent compressed space to discover signatures for classification associated with aging, smoking, and disease. We believe this approach can improve the understanding of complex epigenetic processes and help develop more effective diagnostic and therapeutic strategies.

## INTRODUCTION

Knowledge-driven methods for data analysis in systems medicine involve the use of prior biological understandings to guide and inform the analysis of large data sets. One of the most common of these approaches are network models, which represent biological entities, such as proteins or genes, and their functional relationships as nodes and edges within the interactome network. The interconnected nature of disease genes and their protein products to form disease modules has been exploited by module identification tools that can detect these structures from omics data, validating them by their enrichment in disease-associated SNPs from genome-wide analyses [1] [2]. However, despite curation efforts, human protein-protein interaction (PPI) networks are often incomplete and may not reflect the full complexity of biological systems. They are prone to research biases, as studies may focus on well-known proteins or interactions that are easier to detect [3] [4] [5]. Furthermore, network inference tools often use simplifications in order to construct networks, which can affect the certainty of their predictions [6] [7], and are limited by the coverage and quality of omics data [8]. Incomplete or noisy data can lead to inaccurate networks, reducing their reliability and usefulness. Therefore, there is a need for robust data-driven approaches with the potential to unbiasedly identify novel meaningful signatures.

DNA methylation (DNAm) modifications are well-established biomarkers, due to their capacity to capture long-term environmental effects. DNAm is an ideal modality for data-driven approaches due to its molecular stability, continuously variable nature, and accessibility for large-scale analyses. For instance, it has proven useful in multiple studies of aging, cancer, and other diseases [9]. Changes in the DNAm at specific positions, known as CpG sites, located in genes involved in inflammation and DNA damage responses, have been shown to occur alongside age in a predictable manner [10]. This has allowed researchers to develop algorithms named DNAm clocks, which can accurately estimate chronological age based on an epigenetic profile [11] [12] [13]. Similarly, distinct alterations in DNAm due to tobacco exposure can be used as precise biomarkers of smoking status to assess the effects of smoking habits on gene expression and lung function [14] [15] [16].

Following the same reasoning, while DNAm homeostasis is tightly regulated in healthy individuals, it can become dysregulated in autoimmune diseases like systemic lupus erythematosus (SLE), neurodegenerative disorders, or cancer. In SLE, several genes involved in immune response mechanisms, such as inflammation and antibody production, present abnormal DNAm patterns [17] that have been associated with different degrees of disease activity, severity, and susceptibility [18] [19] [20]. In general,

The emergence of deep learning techniques, based on the concept of artificial neural networks (ANNs), has revolutionized the scientific panorama due to their superb capacity to model complex big data [21] [22] [23]. Autoencoders (AEs) are a type of ANN trained to efficiently compress and reconstruct unlabeled feature sets by learning their internal representations. AEs of different types and configurations have been successfully applied to omic research. For instance, deep undercomplete or variational AE architectures, such as scETM [24], VEGA [25], LDVAE [26], scMVAE [27], scIAE [28], or VASC [29] have been used to analyze single-cell transcriptomic data and identify cellular and gene signatures. Likewise, AEs have also been used to determine disease progression [30] [31], to cluster cancer subtypes [32] [33], to investigate protein variants [34] or to integrate spatial modalities in tissue samples [35]. More recently, the interpretation of the internal embeddings of trained AE models has begun to receive significant attention. For example, modular hierarchical organizational levels of biological interest have been observed within a deep single-cell RNA-seq AE [36] and a deep transcriptomic AE [37]. Further, frameworks such as MethylNet [38] or siVAE [39] demonstrate the potential of interpretable feature embeddings in omic research.

Our workflow seeks to find a compact representation that captures the biological relevance of DNAm data using a minimum number of model parameters. Deep undercomplete AEs are relatively simple to implement and train, while being highly efficient at pattern learning. In addition, hidden representations from deep AEs are deterministic, with inputs being clustered into discrete vectors within a compressed space that is not constrained into a prior distribution. Another benefit of these generic representations is that they are transferable across tasks, meaning that a supervised ANN trained on the latent space can extract task-specific features for any given purpose. This property makes deep AEs particularly attractive for scenarios where large amounts of labeled data may not be available, as the AE can be trained on a much larger, diverse set of data and then be fine-tuned for the particular task at hand.

Here, we propose a novel data-driven approach for the functional analysis of DNAm data using deep AEs with biologically relevant latent embeddings, named network-coherent AEs (NCAEs). We showcase a generally applicable pipeline for the training, selection, and functionalization of deep NCAEs, and demonstrate the potential of their coherent compressed space for the discovery of DNAm signatures and robust signatures for classification. Supervised ANN models trained on these generic transferable features for a variety of tasks performed as accurately or more than gold-standard DNAm-based estimators. In summary, by learning in an unsupervised fashion the inherent structure of DNAm data that incorporates biological knowledge within an interpretable AE design, we are able to uncover meaningful patterns that may not be apparent using conventional methods. Our results show that our data-driven workflow can be used to identify unbiased epigenetic disease and risk factor signatures, and can be generalized to any task where a supervised network is applicable, thus potentially leading to better diagnostic and therapeutic strategies.

## RESULTS

### Deep AEs can accurately reconstruct low dimensional embeddings of methylation data

We searched in an unsupervised manner for a functional data representation that could encompass and exploit the complete feature space, while simultaneously being reduced in its dimension to facilitate predictive modelling and decrease noise. For this aim, we downloaded and pre-processed a multi-tissue compendium of 75,272 human DNAm profiles and metadata from the EWAS (Epigenome-Wide Association Study) Data Hub public repository [40] from Illumina 450K or EPIC arrays. Non-tumor tissue samples (n = 50,623) were included in the model training. Case-control samples from 150 different non-cancer diseases are represented in the collection, obtained from 354 different tissues or cell types (**Fig. 1**). Cancer samples (n = 24,649) were kept as an independent test set to avoid introducing bias into the model. DNAm patterns from tumor tissues are often aberrant [41], so including these samples in the training data could cause the models to learn features that are specific to cancer, rather than more general features that are relevant across a wider range of samples. Non-cancer DNAm profiles were randomly split into training, validation, and test sets balanced for tissue and disease groups (Methods). We trained and evaluated the performance of 24 different undercomplete AE architectures with depths of one to four hidden layers, and widths ranging from 32 to 1,024 hidden nodes in incremental steps of powers of two. Reconstruction performance was measured by the coefficient of determination (R^2^) calculated over the global and local (CpG-specific) variances on the test set. We observed global R^2^ values between 0.978 and 0.993, with decreasing reconstruction error primarily associated with an increase in model width, but not depth, until our maximum tested 1,024 nodes per layer (**Fig. 2a**).

**Fig. 1.**
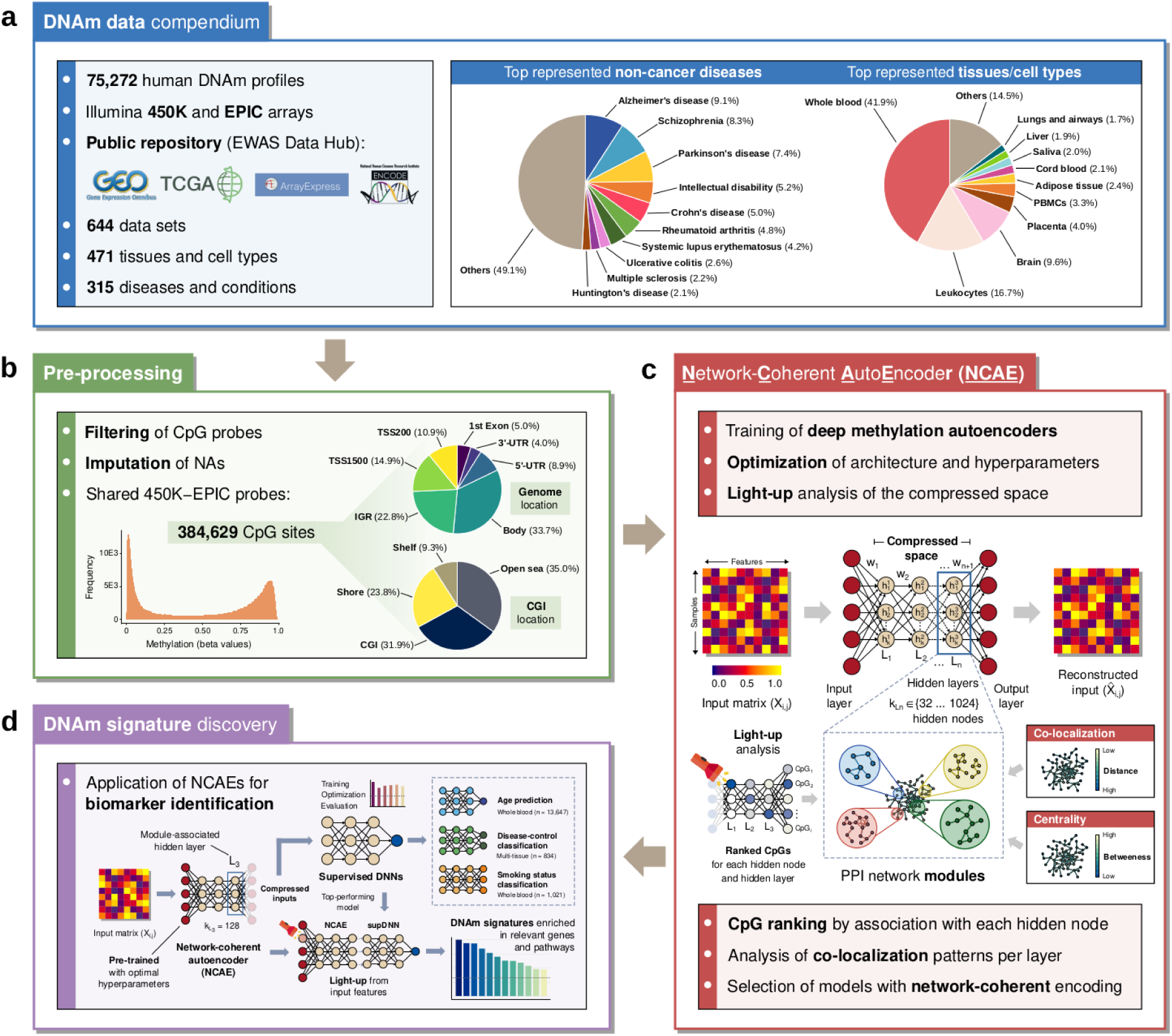
Schematic of the workflow for training and functionalization of network-coherent autoencoders. **a** Summary of DNAm data sets included in the compendium, top represented non-cancer diseases, and tissues and cell types. **b** Pre-processing steps and description of probe set genomic and CpG island (CGI) locations. **c** Training and selection of deep autoencoders based on the network coherence of their latent space (NCAEs). **d** Functionalization of deep NCAE-compressed representations for the identification of DNAm signatures using concatenated task-specific deep supervised neural networks (DNNs).

**Fig. 2.**
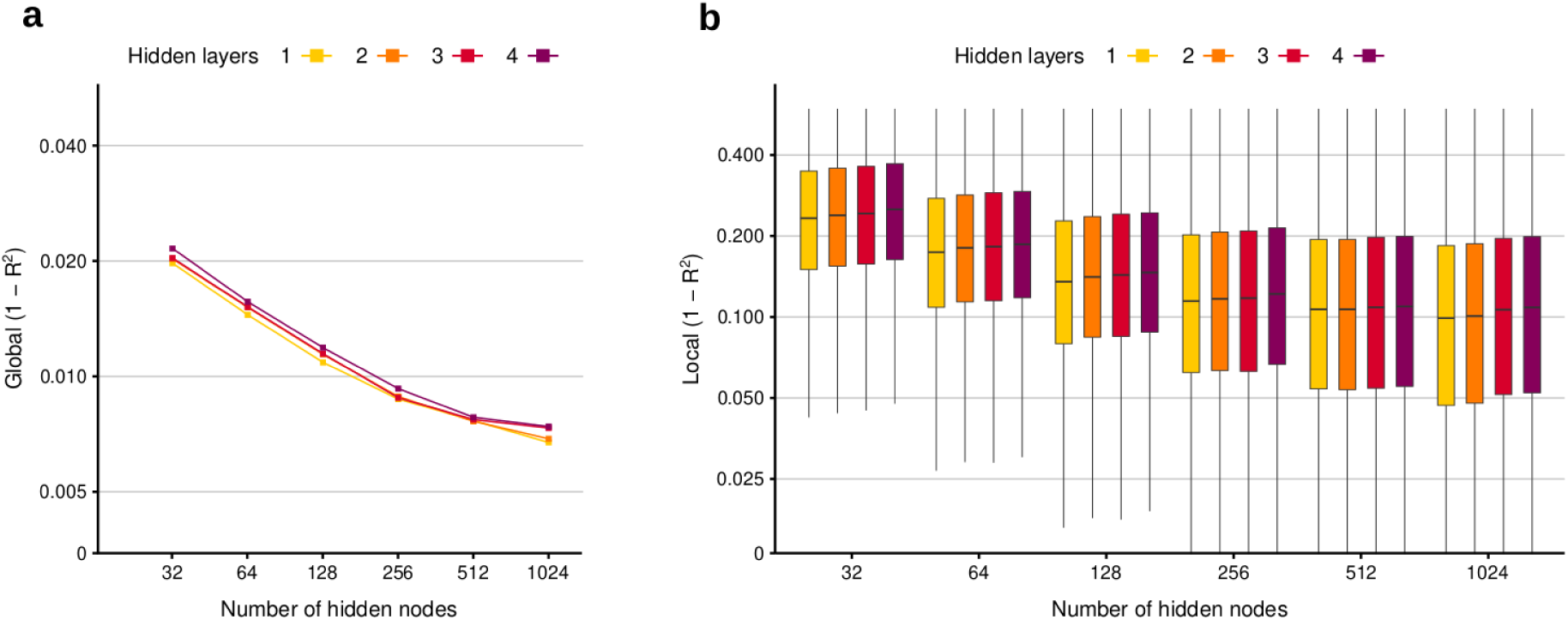
Performance evaluation of methylation autoencoder architectures. Global (**a**) and local (**b**) 1 - coefficient of determination (R^2^) for test set data across AE model configurations, from one to four hidden layers, 32 to 1,024 hidden nodes per layer.

Since some CpGs are prone to have quite stable methylation levels, we next analyzed the proportion of explained variance per CpG (**Fig. 2b**), which showed a similar pattern. AE models of 32 hidden nodes per layer achieved median local R^2^ values between 0.767 and 0.749, while wider configurations such as 512 and 1,024 hidden nodes per layer performed with higher accuracy (median local R^2^ values between 0.901 and 0.891). Specifically, we observed that 85.9% of the variance could be explained for more than half of the CpGs already for models with 128 hidden nodes, corresponding to a 3,000-fold dimensionality reduction (Suppl. File 1). We then investigated whether a fraction of CpGs existed that were consistently well or poorly predicted across the different AE architectures. Notably, we found that 4.7% of the probes (n = 18,118) were systematically below the first quartile (Q1, R^2^ = 0.922) for every AE prediction. These CpGs were more variable than average (mean σ^2^ = 4.63e-2, average mean σ^2^ = 1.65e-2, Wilcoxon P = 0) and included a statistically higher proportion of hypermethylated probes (beta values >= 0.6, 57.1% vs. 50.5%, chi-square P = 1.21e-36) than expected. Conversely, 12.9% of the CpGs (n = 49,463) were consistently reconstructed with a local error above the third quartile (Q3, R^2^ = 0.751). These probes were less variable than average (mean σ^2^ = 4.73e-3, Wilcoxon P = 0) and predominantly hypomethylated (74.0% vs. 43.4%, chi-square P = 0).

### CpGs from the third layer of a deep AE are associated to highly co-localized genes

As this work aimed to find a robust and multi-purpose representation capable of accurately compressing CpG methylation status, we continued to analyze the functional aspects of the above learned representations. To this end, we investigated their large-scale associations to the human protein-protein interaction (PPI) network. Our hypothesis was that a functional representation should cluster CpGs associated to functionally related genes together, thus associating genes with low average distances in the PPI to the same latent variable. We ranked CpGs by their association with each hidden node using light-up analysis and inspected the relationships between the top prioritized CpG-associated genes per hidden layer in terms of their co-localization and centrality. Light-up analyses are a type of perturbation-based forward propagation approach that allow to interpret the non-linear embeddings of an ANN by relating components of the internal layers of the model to its output [37]. Namely, an input vector consisting of a single maximum value of the activation function for a hidden node and null values for all other nodes is propagated through the model layers (Methods). The resulting activation signal can then be used to rank features in the output layer, in this case CpG sites. We repeated the procedure for all hidden nodes and layers, for the different AE architectures.

To determine if the ranked CpGs corresponded to functional structures, we mapped them to their associated genes and analyzed their co-localization in the human PPI network defined by STRING v11 [42] high-confidence interactions (score > 0.7). We calculated the harmonic average distance (HAD) between the gene lists obtained for every model and layer, and compared it with the average HAD within the PPI network (HAD = 3.48). Gene sets with a low mean HAD have a higher degree of co-localization, since they are situated closer within the interactome. The analysis revealed that top ranked genes were more co-localized than expected (Wilcoxon rank sum P = 0), and this decrease in mean HAD was higher the further into the AE layers the light-up signal was propagated from (Suppl. File 2). The effect reached a plateau at the third layer of the four-layered AE models, with the top genes associated with the fourth layer showing similar HAD. Top genes from the third layer of three-layered AEs presented the lowest HAD, especially notable for models with widths of 128 and 256 hidden nodes (**Fig. 3c**). By contrast, top genes from the first and second hidden layers only showed a weak co-localization increase, while still being below the average HAD (**Fig. 3a, b**). This effect was stronger for the top 100 to 400 ranked genes, vanishing progressively through the rest of the ranking. In order to further understand the association, we also analyzed the betweenness centrality of the associated CpGs, which yielded a similar outcome (**Fig 3d**). The most central genes associated most with the latent variables of the third layer of the three-layered AEs with 128 to 512 nodes, whereas the effect was lower for other layers and depths (Suppl. File 2).

**Fig. 3.**
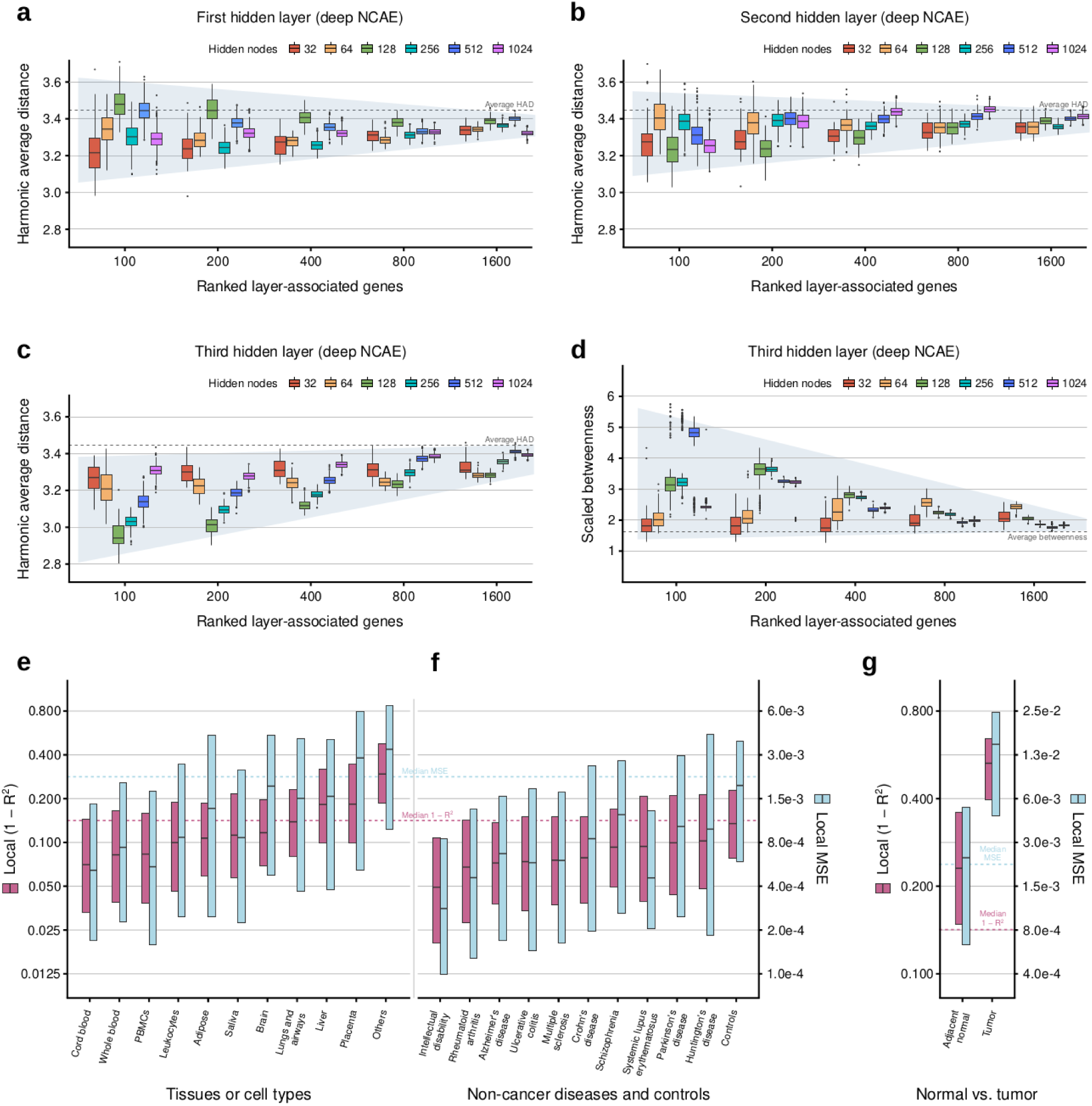
The third hidden layer of a deep methylation AE shows gene co-localization patterns in the human interactome. Harmonic average distances in the human interactome of the top 1,600 ranked genes associated to the first (**a**), second (**b**), and third (**c**) hidden layers of the deep NCAE with three hidden layers and 128 hidden nodes per layer. **d** Betweenness centrality of genes associated to the third hidden layer of the NCAE. Local reconstruction performance (1 - R^2^) of the NCAE on test set top represented tissues and cell types (**e**), top represented diseases, plus controls (**f**) and tumor and adjacent normal tissue samples (**g**).

In summary, we observed that the corresponding latent variable representations differ substantially in terms of their relation to the PPI network. Specifically, CpGs associated with the third layer of the three-layered deep AE of 128 hidden nodes showed the strongest co-localization pattern (HAD = 2.9) and above-average centrality within the human PPI network, while ranked CpGs from other architectures and layers had weaker patterns and associated with disparate areas of the interactome. We believe that such a functional and also substantially reduced representation is highly valuable for transfer learning and multi-purpose training of neural networks for research questions for which fewer labeled samples are available, and that this functional relation enables the identification of new CpG associations in a data-driven manner. Therefore, we subsequently focused our analysis on the three-layered deep AE of 128 nodes, which will henceforth be referred to as network-coherent AE (NCAE). We replicated the analyses in sparse AEs and variational AEs with architectures equivalent to the NCAE (Methods), but we did not found a significant improvement in co-localization (Suppl. File 3).

### The deep NCAE reconstructed patients of most common non-cancer diseases and tissues as good as controls

Next, we tested if the high-level compression of the NCAE was biased towards certain tissues or diseases. To do so, we calculated the CpG-wise R^2^ across test set samples from the ten most frequent tissues or cell types and the ten most frequent non-cancer diseases, and healthy controls. Interestingly, well-explained tissues were related to circulating blood cells, such as cord blood, whole blood, PBMCs; whereas localized tissues, such as liver, brain, or placenta, had lower R^2^ (**Fig. 3e**). We found no significant correlation (Pearson *r* = 0.36, P = 0.304) between the NCAE performance and the proportion of samples per tissue in the training set. The top represented non-cancer diseases showed median R^2^ values between 0.951 and 0.898, whereas controls were predicted with R^2^ = 0.865 (**Fig. 3f**), likely due to a higher variability across healthy individuals. Again, the NCAE performance and the proportion of samples per disease in the training set were not significantly correlated (Pearson *r* = -0.01, P = 0.978). Lastly, we analyzed the NCAE performance on the left-out cancer samples (n = 24,649) and their respective test set adjacent normal tissue samples (**Fig. 3g**). Tumor samples were explained poorly, achieving only R^2^ = 0.471, compared to R^2^ = 0.769 for adjacent normal tissue. Presumably, the NCAE performance on cancer samples would improve if included in the training set. However, methylation features unique to tumor DNAm profiles could result in the formation of clusters on the latent space occupied by cancer samples and enriched on unusual patterns, potentially compromising the ability of the model to learn biologically relevant features. Overall, these results indicate that abnormal tumor-associated methylation patterns need to be considered for the accurate modelling of cancers, but also that the latent representation can capture DNAm patterns of common diseases within relatively low error margins while achieving a proficient accuracy in key cell types for biomarker identification. This led us to further examine how well the latent variables could be repurposed for phenotypic predictions associated to known epigenetic signatures.

### An NCAE-Age model accurately estimates chronological age and identifies relevant aging DNAm signatures

A common task for which DNAm data modelling is highly suitable is the estimation of age using DNAm clocks. Two of the most popular are the hallmark clocks of Horvath [11] and Hannum [12], which can predict chronological age with high accuracy (reported test R^2^ = 0.922, R^2^ = 0.963, respectively). Several studies have suggested an increase in predicted age in association to several diseases, the so-called “accelerated aging” [43] [44] [45] [46]. Remarkably, these two clocks use only 353 and 71 CpGs, respectively, although aging-related processes are likely be spread across many more DNA regions. Thus, a broader and more robust DNAm clock could potentially serve better to understand aging. For this purpose, we used the learned compressed representation of 128 latent variables from the NCAE to train a deep supervised neural network (NCAE-Age) to predict chronological age (Methods) and compared its performance with Horvath and Hannum DNAm clocks.

First, we fed the NCAE a total of 13,647 whole blood DNAm samples from healthy individuals with ages between 0 and 112 years (mean = 40.5 ± 23.6 years, Suppl. File 4) from 67 data sets, to extract the latent space embeddings and train the NCAE-Age model. Notably, the NCAE-Age achieved highly accurate results on the test set (n = 2,729, R^2^ = 0.965), followed by Horvath (R^2^ = 0.936) and Hannum clocks (R^2^ = 0.929) (**Fig. 4a, b**). Restricting the analysis to test samples in the age range of 19-101 years to match Hannum’s training set did not improve its prediction accuracy (R^2^ = 0.859). The error of the NCAE-Age model was lower than the linear estimators as well (**Fig. 4b**). To better assess the performance of the models across specific age ranges, we binned the test samples into four biologically meaningful age groups (Newborn-Adolescent: 0-18 years, n = 746; Adult: 19-44 years, n = 638; Middle-age: 45-64 years, n = 901; and Aged: ≥65 years, n = 444), according to the Medical Subject Headings (MeSH) criteria. The NCAE-Age estimates were more accurate than the DNAm clocks in all four bins (**Fig. 4c**). All models performed well (R^2^ > 0.650) on the age groups between 0 and 44 years, with a pronounced decrease of accuracy above that threshold, especially for Horvath clock in the Aged group (NCAE-Age R^2^ = 0.523, Hannum R^2^ = 0.510, Horvath R^2^ = 0.333). We observed a tendency of the Hannum clock to underestimate sample ages above 40-50 years, which substantially increased the absolute error of its predictions in the corresponding age groups, up to 14.28 years in the Aged bin. In general, the performance of the NCAE-Age model trained with the compressed feature embeddings was comparable or superior to Horvath and Hannum DNAm clocks.

**Fig. 4.**
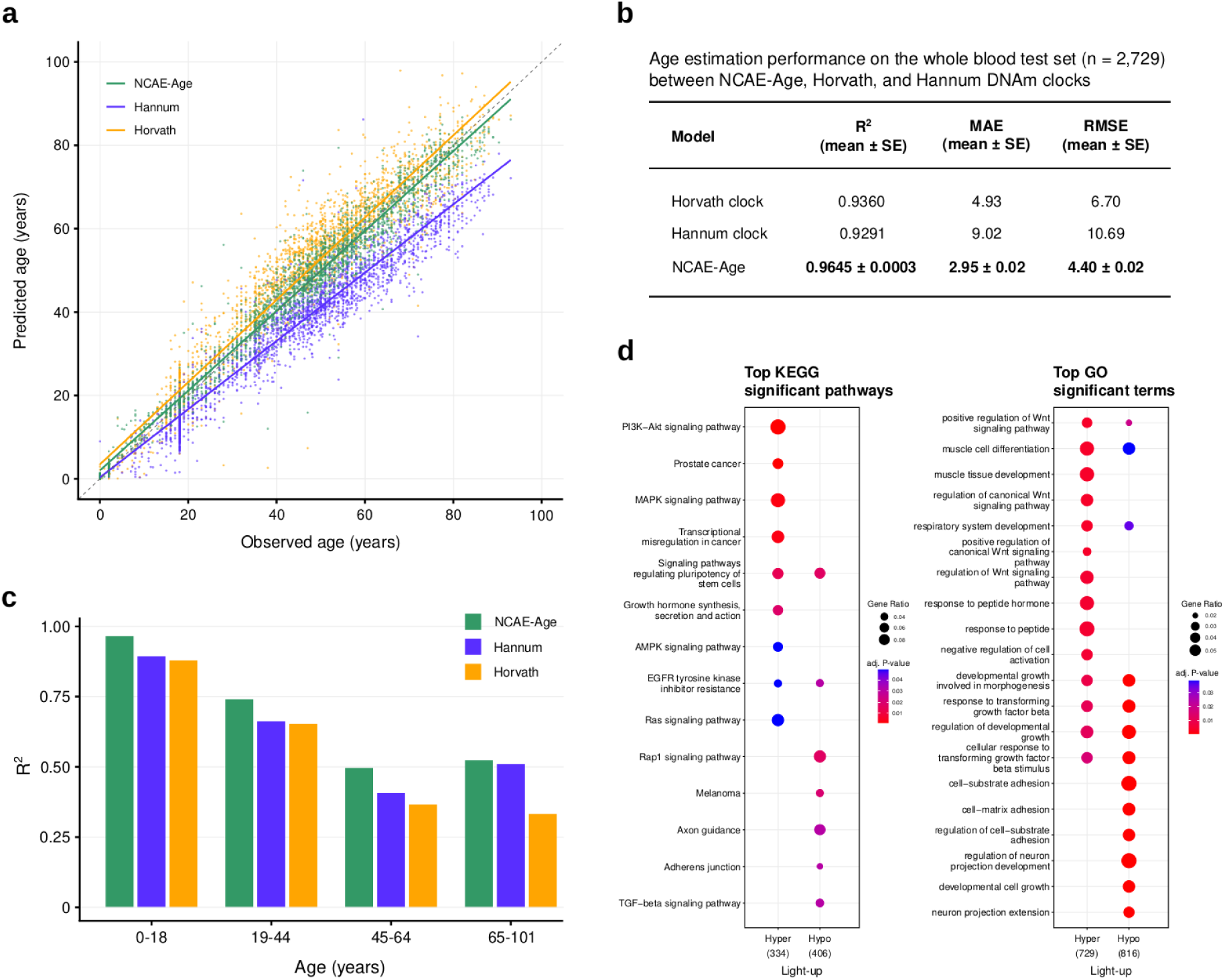
A non-linear DNAm age estimator (NCAE-Age) trained on whole blood NCAEcompressed inputs can predict age more accurately than hallmark DNAm clocks. **a** Comparison of true and predicted ages of test set samples (n = 2,729) as estimated by NCAE-Age, Horvath, and Hannum DNAm clocks. Prediction performance of the DNAm estimators on the full test set (**b**) and on the binned samples by age group (0-18: Newborn-Adolescent, 19-44: Adult, 45-64: Middle-age, 65-101: Aged) (**c**). d Top significantly overrepresented KEGG pathways and GO terms for the top ranked CpG-associated genes after the light-up analysis.

To assess the leverage of particular CpG sites in the NCAE-Age estimates and determine the model’s ability to prioritize CpGs associated with genes involved in DNAm predictors and known age-related mechanisms, we applied a light-up analysis from the input of the NCAE (Methods). Briefly, we forward-propagated a recursive perturbation signal per input CpG through the concatenated hidden layers of the AE and supervised DNN, and across a tissue-specific methylation profile. The effect of each perturbation in the model output, measured as changes in the predicted age, can be used to rank CpGs by the relevance of their contribution to the model. We then tested whether the resulting lists of top ranked CpG-associated genes were enriched in nine chronological and five biological DNAm clock genes (n = 1,328, Suppl. File 5) [47], developed mainly using blood or saliva as DNA source. Reassuringly, we found that genes associated with the top 1,000 CpGs identified by hyper- or hypo-perturbation significantly overlapped the combined age clock gene list (P = 3.49e-13, OR = 2.23; and P = 2.27e-5, OR = 1.59, respectively). In particular, the top ranked CpG list obtained by the hyper-methylation light-up analysis (Suppl. File 6) was significantly enriched in every DNAm clock list containing more than four genes (ten DNAm clocks, P = 1.71e-8 to 0.04, OR = 1.59 to 80.02, Suppl. File 7).

Next, we investigated if the prioritized genes are known to participate in biological pathways linked to functional processes in human aging. The KEGG gene enrichment analysis of the top ranked lists revealed a significant (adj. P < 0.05) overrepresentation of signaling pathways (PI3K-Akt, MAPK, AMPK, Ras, Rap1, TGF-beta; adjusted P = 1.19e-3 to 0.049), and cancer-related pathways (transcriptional misregulation in cancer, prostate cancer, melanoma; adj. P = 1.19e-3 to 1.99e-2), among others such as axon guidance and adherens junction (adj. P = 3.08e-2, both). Furthermore, the list obtained by input hyper-perturbation presented a significant enrichment in Disease Ontology (DO) terms linked with multiple types of cancer (adj. P = 5.64e-4 to 4.60e-2). Interestingly, both lists were highly enriched in Gene Ontology (GO) terms associated with responses to transforming growth factor (TGF) beta (adj. P = 2.52e-5 to 1.04e-2) and positive regulation of the Wnt signaling pathway (adj. P = 3.73e-3 to 1.93e-2) (**Fig. 4d**).

### An NCAE-Smoke model determines smoking status and defines associated smoking DNAm signatures

Besides aging, another common use case within DNAm research concerns the modelling of alterations in methylation patterns due to tobacco smoking, for which clear DNA signatures have been identified, even long after cessation. We hypothesized that these changes in the DNAm landscape caused by smoking may be imprinted on the deep NCAE compressed representations, and thus epigenetic signatures for smoking can be derived from them.

We retrieved 1,021 whole blood DNAm samples from individuals enrolled as controls in five studies with available self-reported smoking status information (Suppl. File 4). The selection of whole blood was motivated by the abundance of related samples in epidemiological studies, as well as its relevance in the identification of biomarkers for smoking-related disorders. We obtained the compressed embedding using the NCAE and trained a deep supervised neural network (NCAE-Smoke) for multi-class classification of samples as belonging to a current (n = 408), former (n = 185), or never smoking individual (n = 428), where 20% of samples in each group were used as test set (Methods). We observed the best performance on the test set in an L1-regularized three-layered NCAE-Smoke with leaky ReLU activation on the hidden layers and softmax activation on the output layer (**Fig. 5a**, AUC_current_ = 0.87, AUC_former_ = 0.91, AUC_never_ = 0.69, average AUC = 0.80).

**Fig. 5.**
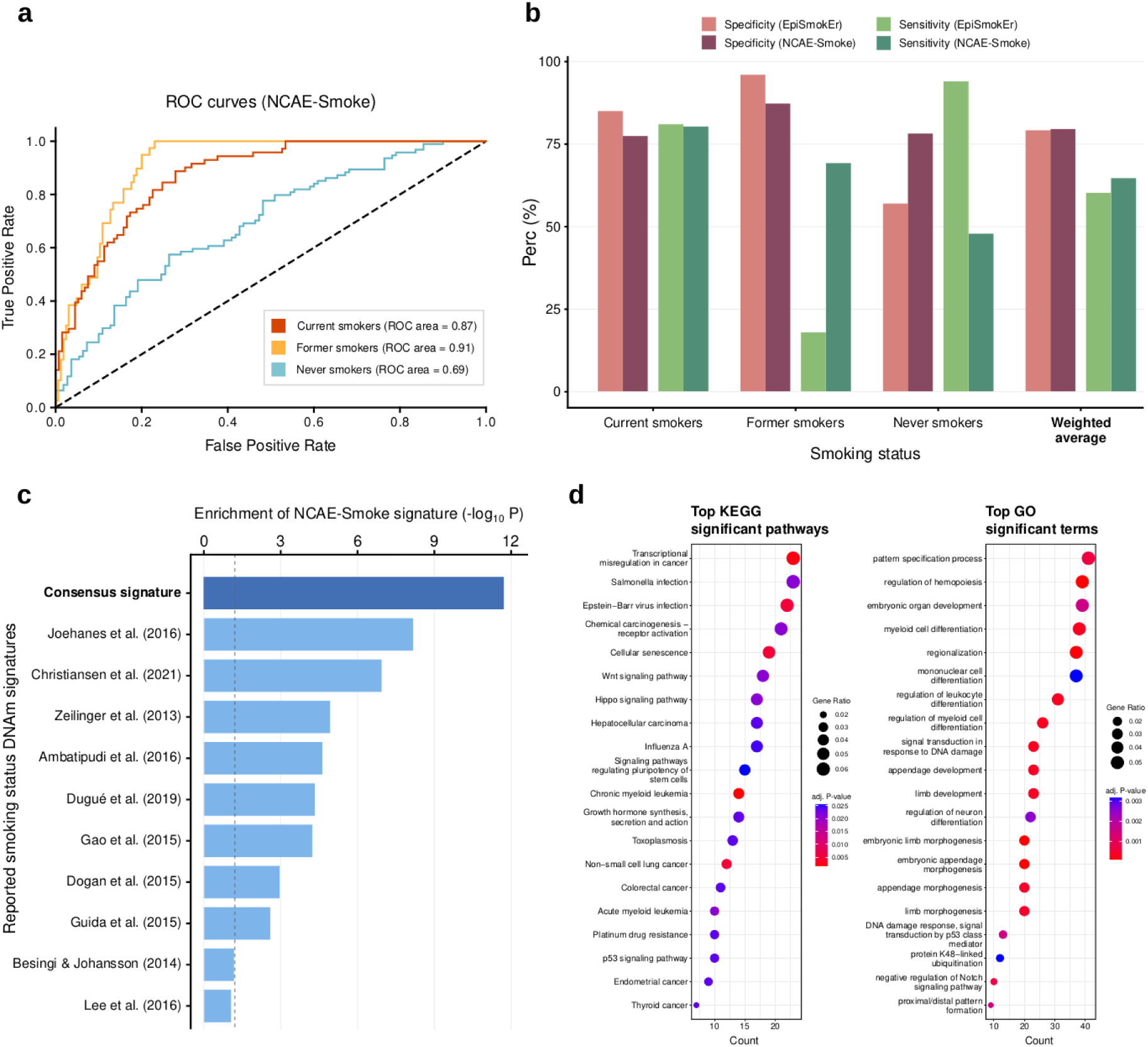
A non-linear DNAm smoking status classifier (NCAE-Smoke) trained on NCAE-compressed inputs accurately separates current, former, and never smokers. **a** AUROC curves for NCAE-Smoke classification performance of smoking status classes. **b** Comparison between NCAE-Smoke and EpiSmokEr reported average per class specificity and sensitivity on the test set, and weighted average by sample size across classes. **c** Significance of the enrichment of top ranked CpG-associated genes from the NCAE-Smoke, for current vs. never smokers, in reported DNAm signatures for smoking status. The consensus signature includes all genes appearing in at least two of these studies. **d** Most significantly overrepresented KEGG pathways and GO terms for the top ranked NCAE-Smoke CpGs for current vs. never smokers.

Next, we evaluated the classification performance of the NCAE-Smoke in comparison with other available DNAm-based smoking status predictor trained on whole blood data. EpiSmokEr [48] uses 121 CpG coefficients identified via a LASSO-penalized generalized linear model, plus a sex and an intercept coefficient, to determine smoking status on the same three categories (current, former, never smoker). Compared with the reported performance of EpiSmokEr on the test set data (**Fig. 5b**), both models were able to correctly categorize current smokers (EpiSmokEr Spec = 85%, Sens = 81%; NCAE-Smoke Spec = 77%, Sens = 80%), as well as reliably ruling out samples that are not former smokers (EpiSmokEr Spec = 96%, NCAE-Smoke Spec = 87%). However, NCAE-Smoke achieved a higher true positive rate for former smokers (Sens = 69%) than EpiSmokEr (Sens = 18%). Regarding never smokers, NCAE-Smoke performed better in terms of specificity (77% vs. 57% for EpiSmokEr), whereas EpiSmokEr had a higher sensitivity (94% vs. 48% for NCAE-Smoke). On average, across the different smoking status classes, the performance of NCAE-Smoke was similar or above EpiSmokEr reported values (NCAE-Smoke Spec = 79%, EpiSmokEr Spec = 79%; NCAE-Smoke Sens = 65%, EpiSmokEr Sens = 60%).

Therefore, we fixed the model architecture and hyperparameters, trained a NCAE-Smoke on the complete data, and performed an input light-up analysis. Hence, we retrieved a list of CpGs prioritized by differences between the “never smoker” and “current smoker” classes, selecting the genes associated to the top 1,000 CpGs as the NCAE-based DNAm smoking signature (Suppl. File 6). To determine whether these top ranked genes were indeed linked to altered methylation due to ongoing smoking, we evaluated the significance of their overlap with ten available DNAm signatures for smoking status (n = 95 to 3,978 CpGs, Suppl. File 8). We observed that CpG-associated genes from eight out of ten reported DNAm signatures were significantly overrepresented (Fisher’s exact P = 6.60e-9 to 2.48e-3, OR = 2.61 to 8.18) in the NCAE-based ranked gene list (**Fig. 5c**). Moreover, a consensus DNAm signature including genes that appear at least twice across the reported smoking signatures (n = 1,462) showed the most significant overlap with the NCAE signature (Fisher’s exact P = 1.87e-12, OR = 2.09).

Examining the biological context of this DNAm signature for never smokers vs. current smokers (**Fig. 5d**), we found that it was enriched in KEGG pathways associated with cancer, the most significant ones being myeloid leukemia (adj. P = 1.57e-3 to 2.11e-2), non-small cell lung cancer (adj. P = 5.22e-3), transcriptional misregulation in cancer (adj. P = 2.18e-3), p53 signaling pathway (adj. P = 2.38e-2), and chemical carcinogenesis receptor activation (adj. P = 2.11e-2). Further, the signature was also enriched in GO terms linked to DNA damage, such as signal transduction in response to DNA damage (adj. P = 3.03e-4), DNA damage response, signal transduction by p53 class mediator (adj. P = 1.55e-3 to 2.85e-2), mitotic DNA damage checkpoint signaling (adj. P = 2.47e-2). Other significantly overrepresented terms were embryonic morphogenetic processes (adj. P = 8.41e-5 to 4.33e-2) and regulation of myeloid cell differentiation (adj. P = 2.71e-4 to 4.58e-2).

### An NCAE-SLE model for tissue-specific disease biomarker discovery using latent space features

Thus far, the latent AE embeddings of the DNAm data have been shown to organize in an unsupervised manner into module-like patterns that can encompass age and smoking status information. We further hypothesized that epigenetic disease biomarkers could also be detected in the NCAE-compressed feature space. SLE is considered as the prototypical autoimmune disorder, coupled with the relevance of DNAm alterations in immune cells of SLE patients, makes it an ideal candidate for this analysis.

We compiled six available data sets containing DNAm profiles from SLE patients and healthy controls (n = 834, Suppl. File 4) across 11 tissues or cell types. Then, we trained a multi-tissue deep supervised neural network (NCAE-SLE) for SLE patient-control classification. We used the NCAE latent embeddings, propagated in an L1-regularized three-layered NCAE-SLE, to differentiate SLE cases from healthy individuals on the multi-tissue test set (n = 167, accuracy = 0.78, AUC = 0.89). Evaluating the classification performance per tissue or cell type (**Fig. 6a**), the DNN showed the best results for DNAm samples from T cells (n = 76, acc = 0.87, AUC = 0.95), PBMCs (n = 13, acc = 0.77, AUC = 0.93) and monocytes (n = 21, acc = 0.86, AUC = 0.85). Less represented cell types, such as neutrophils (n = 7) and granulocytes (n = 6) achieved lower accuracy (AUC = 0.50, AUC = 0.67, respectively).

**Fig. 6.**
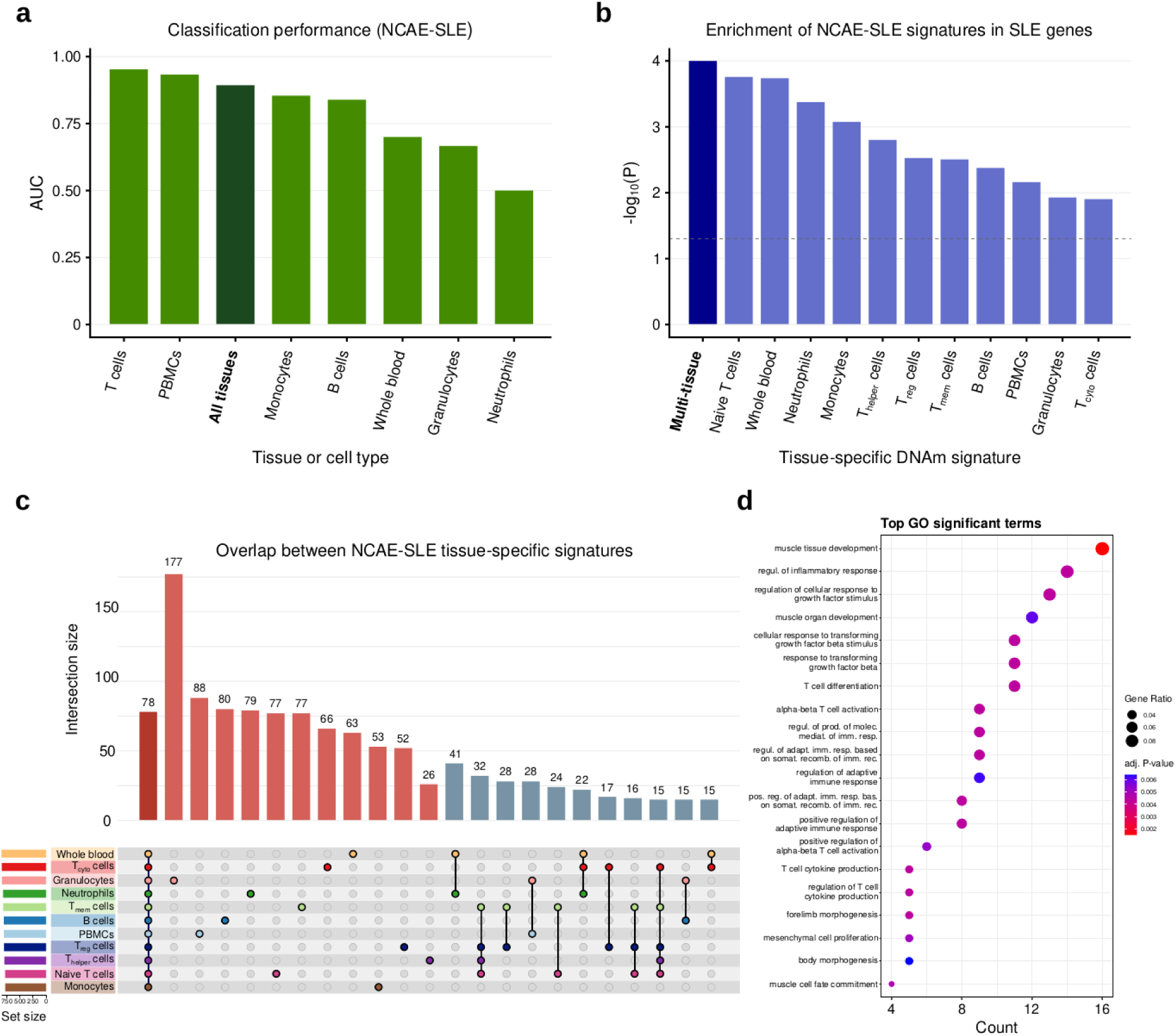
A non-linear DNAm multi-tissue SLE patient-control classifier (NCAE-SLE) trained on NCAE-compressed inputs can be used to obtain disease-associated methylation signatures. **a** AUC values for NCAE-SLE test set classification performance across tissues and cell types. **b** Significance of the enrichment in SLE-associated genes from DisGeNET of top ranked CpG-associated genes by tissue and cell type from the trained NCAE-SLE. Multi-tissue refers to the top 1,000 most frequent genes across the single-tissue NCAE-SLE rankings. **c** Overlap between NCAE-SLE CpG-associated genes from the top light-up ranked 1,000 CpGs per tissue. **d** Top significantly enriched GO terms for the multitissue SLE DNAm signature.

Once the model configuration was optimized, we trained a NCAE-SLE with identical architecture and hyperparameters on the entire data (Methods) and performed a light-up analysis from the AE input to identify whether CpGs prioritized by the model are linked to known SLE-associated genes retrieved from DisGeNET v7.0 [49]. Since 11 tissues and cell types were present in the SLE data set compilation, light-up was applied on different average DNAm profiles for each of them. Tissue- and condition-specific DNAm profiles were recursively perturbed to obtain lists of CpGs ranked by their association to SLE according to the trained NCAE-SLE model. The effect of CpG perturbations in the output of the model was measured as the absolute change in the predicted probability of a DNAm profile to be assigned as a SLE case or control, depending on the profile.

To better understand the model capacity to detect disease-linked markers with fine-grained resolution, we computed the cumulative enrichment in SLE-associated genes for the genes associated to the top 1,000 light-up ranked CpGs per tissue (Suppl. File 6). We assessed the significance of the observed enrichments using a permutation test to estimate a P-value (n = 1e4 permutations). We found significant enrichments in SLE genes in the hyper-methylation light-up CpG lists for all the tissue-specific sets (**Fig. 6b**). CpG lists obtained from naive T cells and whole blood were particularly strongly enriched in disease genes (P = 1.75e-4, P = 1.83e-4, respectively). Furthermore, we observed that a multi-tissue list containing the first 1,000 genes by frequency across the top CpGs per tissue or cell type achieved the highest possible enrichment for the number of permutations (P = 1e-4). The subset of genes (n = 184) appearing at least once per single-tissue NCAE-SLE signatures was also highly enriched in DisGeNET genes for SLE (Fisher’s exact test P = 2.70e-4, OR = 2.00). These results suggest that the NCAE-SLE model can discern tissue-specific and multi-tissue gene-disease associations from the NCAE-compressed features, from which relevant candidate biomarker lists can be retrieved.

We then performed gene enrichment analyses to identify the biological context of the ranked gene lists in relation to known SLE-associated pathological mechanisms. Considering the tissue-specific light-up results (**Fig. 6c**), we found multiple hits for significant (adj. P > 0.05) enrichments in KEGG pathways linked to transcriptional misregulation in cancer, Epstein-Barr virus infection, Th17 cell differentiation, antigen processing and presentation, and the Hippo and FoxO signaling pathways, among others. In like manner, they were highly enriched in GO terms such as antigen processing and presentation, morphogenesis-related terms, and T cell mediated immunity and differentiation. With regard to the multi-tissue SLE DNAm signature, we observed strong enrichments in GO terms linked with muscle tissue development (adj. P = 1.50e-3 to 4.74e-2) and morphogenetic processes (adj. P = 4.41e-3 to 4.33e-2), immunoregulatory processes related with the adaptive immune response (adj. P = 4.41e-3 to 1.68e-2), TGF-beta (adj. P = 4.41e-3 to 3.76e-2) and T cell cytokine production (adj. P = 4.41e-3 to 4.97e-2), and mesenchymal cell proliferation (adj. P = 5.06e-3 to 3.94e-2) (**Fig. 6d**).

## DISCUSSION

Here, we introduced a deep learning workflow for the identification of network-coherent AEs (NCAEs), which encode a biologically meaningful latent space that can be used for DNAm signature discovery. Assessing the performance of an autoencoder is a non-trivial task, since simply comparing the reconstructed data to the original input may not be sufficient to determine the usefulness of the learned representation. Our study aimed at demonstrating whether the interpretable embeddings of a deep AE trained on large DNAm data can capture complex, non-linear relationships of biological relevance that could be used for selecting a best-performing autoencoder, from which novel unbiased epigenetic signatures can then be detected. We examined multiple architectures (two to four hidden layers, 32 to 1,024 hidden nodes per layer) and hyperparameter configurations of deep AEs trained on a multi-tissue DNAm compendium to determine a configuration that balanced reconstruction performance and coherence with the human protein interactome within its encoding. Wider AE models explained DNAm data better than deeper ones, with the slight decrease in performance for subsequent hidden layers probably due to the difficulties to train a more complex network architecture and the higher risks of information loss.

From the trained models, the latent space analysis via light-up revealed that CpGs associated to genes with a high co-localization in the human PPI network were increasingly prioritized alongside model depth, until the third layer. We selected the deep methylation AE with three hidden layers and 128 hidden nodes per layer as our NCAE. The observed gene localization gradient suggests that different hidden layers encode different biological signals, as shown previously for a deep transcriptomic AE [37]. Further, the late emergence of the co-localization signal may indicate that processes of a higher order are modelled before the association to the PPI is decoded, particularly since an increase in central genes was observed in parallel. We showed that the latent embeddings of the pre-trained NCAE can be functionalized for transfer learning-based candidate biomarker discovery, mapping DNAm data to the biologically relevant compressed space, before feeding the new feature set to a concatenated supervised deep neural network. Lastly, we applied feature-wise input light-up to rank CpGs by their association to the training objective of the NCAE-DNN, thus obtaining task-specific epigenetic signatures.

We validated this approach on three use cases: age estimation, smoking status, and SLE patient-control classification. The NCAE-Age performed as well or above Horvath and Hannum DNAm clocks across every age group. Genes linked to CpGs from a list of DNAm age estimators were significantly overrepresented in the NCAE-Age DNAm signature, which was also associated to pathways known to regulate key aging mechanisms. Inhibition of signaling via PI3K-Akt leads to telomeric DNA damage [50], while its activation may have protective effects against age-modulated neurodegenerative diseases [51]. Similarly, dysregulations within the Wnt signaling pathway have been linked to aging-associated disorders [52], while active TGF-β signaling is connected to accelerated organ senescence [53]. With regard to the NCAE-Smoke classifier, its performance was on pair with the smoking status predictor EpiSmokEr. The resulting top ranked genes were validated against other existing DNAm signatures for smoking status, and they were strongly enriched in pathways related to smoking effects, such as DNA damage response mechanisms, p53 signaling, chemical carcinogenesis, or non-small cell lung cancer [54] [55] [56]. Thirdly, the multi-tissue NCAE-SLE classifier allowed to obtain tissue-specific SLE DNAm signatures, validated using DisGeNET disease-gene associations. These and the combined multi-tissue SLE signature were enriched in processes related to the course of autoimmune diseases, e.g., adaptive immune response regulation, cytokine production by T cells, cellular responses to TGF-β [57] [58] [59].

Comprehensibly, future work may involve efforts to further develop our workflow, such as including paired data from other modalities across a wider range of diseases or environmental factors, such as diet or toxin exposure. Also, the simultaneous modelling of different conditions, like comorbidities or smoking-induced aging, via multi-objective supervised NCAE-DNNs, and the use of more complex models such as graph convolutional networks (GCNs) may improve the relevance of the prioritized CpGs. Overall, our findings suggest that deep NCAEs may be a valuable tool for identifying condition-specific DNAm signatures. We have shown that deep methylation AEs can effciently compress human DNAm data into a co-localized latent space, and that the compressed representation from the deep AE with highest biological coherence (network-coherent autoencoder, NCAE) can be functionalized for effcient ANN training and CpG prioritization. DNAm signature genes from trained NCAE-DNNs are significantly enriched in biological processes related to the condition of interest, resulting in an efficient data-driven biomarker discovery workflow applicable to Illumina 450K and EPIC array data.

## MATERIALS AND METHODS

### Data pre-processing

Human DNA methylation profiles and metadata (n = 75,326) were downloaded from the EWAS (Epigenome-Wide Association Study) Data Hub public repository (https://ngdc.cncb.ac.cn/ewas/datahub, accessed on 2021/01/25). Sources for this database include Gene Expression Omnibus (GEO), ArrayExpress, The Cancer Genome Atlas (TCGA), and Encyclopedia of DNA Elements (ENCODE). Methylation profiles from EWAS Data Hub were generated by Illumina Infinium HumanMethylation450 or MethylationEPIC arrays and were normalized and corrected for batch effects using Gaussian Mixture Quantile Normalization (GMQN) [60]. After sample quality control, 75,272 samples were kept.

Additional filtering was performed using the ChAMP package (version 2.26.0) for the R programming environment (version 4.2.1). Non-CpG probes, probes related to SNPs, multi-hit probes, and probes located on the X or Y chromosomes were filtered out. Beta (β) values for NA probes were imputed using the kNN method (k = 10) from the bnstruct R package (version 1.0.12). After filtering and excluding probes that are not shared by both Illumina 450K and MethylationEPIC arrays, a total of 384,629 CpG sites were left (Suppl. File 9).

Samples obtained from tissues with known aberrant methylation patterns (e.g., cancer-related samples) were excluded from the model training. Therefore, the pre-processed methylation data consisted in a beta-values matrix of 50,623 methylation profiles (Suppl. File 3) by 384,629 CpG sites.

### Design and training of neural network models

Artificial neural network models were trained using Keras 2.4.3 library with TensorFlow 2.4.0 and TensorFlow-GPU 2.2.0 backend, implemented for Python 3.8.10. The normalized DNAm matrix was used to train and evaluate the deep methylation AEs (DMAEs), sparse AE (spDMAEs), and methylation variational AE (MVAEs) models, with a training/validation/test split ratio of 64:16:20, respectively, balanced for tissue and sample group proportions using multivariate stratified sampling. To inspect the impact of the number of hidden nodes on reconstruction performance, we chose to use continuous-width AE models, following the rationale of Sanjiv et al. (2020) [10.1038/s41467-020-14666-6]. Prior to training, hyperparameter optimization was performed using balanced sample subsets of 10-20% of the original population. The optimal configuration used the Adam optimizer to minimize the mean squared error, with learning rate = 9.0e-5, β1 = 0.9, β2 = 0.999, ε = 1e-8, decay = 1e-6. We selected the leaky rectified linear unit (leaky ReLU, α = 0.3) function as hidden layer activation function, and the sigmoid function for the output layer. Dropout regularization did not improve performance and was therefore not applied in the final model. To avoid overfitting, models were trained using early stopping, with a patience of 10 epochs. The batch size for training was 128. The architecture and hyperparameters of the spDMAE and MVAE were matched to that of the network-coherent DMAE of 3 hidden layers and 128 hidden nodes per layer. Thus, the MVAE had 3 dense encoding layers of 128 nodes per layer and a latent Gaussian space of size 2, plus a dense decoding layer. The spDMAE included an L1 activity regularizer constraint = 1e-3 to induce sparsity in each hidden layer.

Supervised DNN models trained on the deep NCAE latent representations were designed to be feed-forward, fully connected neural networks with an input layer of the same dimension as the feature embeddings from the NCAE (128 nodes). To better take advantage of the number of available samples, the DNN training strategy applied was as follows: a) split the NCAE feature embeddings into training, validation, and held-out test sets (64:16:20 ratio) to explore model architectures and hyperparameter configurations that minimized loss, b) once an optimal model configuration is found, train a DNN with identical architecture and hyperparameters on the full training and validation sets (80:20). We used the ADAM optimizer with the same hyperparameters as previously to minimize the MSE (NCAE-Age), categorical cross-entropy (NCAE-Smoke), or binary cross-entropy (NCAE-SLE). The multi-tissue and whole blood NCAE-Age models used ReLU as hidden layer activation function, and leaky ReLU as output layer function. For NCAE-SLE and NCAE-Smoke, we opted for leaky ReLU in the hidden layers, and sigmoid or softmax function in the output layer, respectively. In all cases, L1 kernel regularization was applied on the third hidden layer to prevent overfitting, with a scale factor λ = 0.01. NCAE-DNNs were trained with early stopping (patience = 1e3). The batch sizes used were 1,024 for NCAE-Age, and 256 for NCAE-Smoke and NCAE-SLE. To determine whether sample age and gender could be relevant DNN covariates for smoking status prediction, we trained three additional NCAE-Smoke models using the sample embeddings plus each covariate or both as inputs of the DNN. However, we did not observe an increase in classification recall. Thus, age and gender were not used as DNN covariates.

### Performance evaluation

Reconstruction, regression, and classification performance of ANN models were assessed using metrics from the Python library scikit-learn 0.24.2. Test set local (CpG-wise) reconstruction performance for AEs was measured using the coefficient of determination (R^2^) computed as:

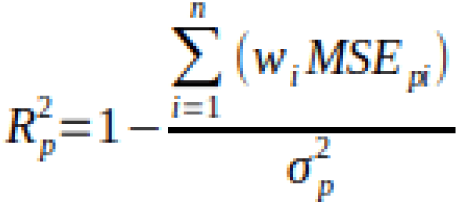

where p, i, wi, and σi2 correspond to the pth CpG probe, the ith data set, the weight (number of samples divided by the total number of samples in the test set) of the ith data set, and the variance of the pth CpG probe.

### Light-up analysis from hidden nodes

To determine the association of AE hidden layers with input CpGs, we retrieved the output layer activations computed from the recursive light-up activation of each hidden node in every layer. That is, we forward-propagated an activation vector *x^a^* consisting of the maximum activation value for a single hidden node *h* of a hidden layer *k*, while keeping the rest of the nodes at the minimum activation. Maximum (1) and minimum (α) activation values used corresponded to the derivative of the AE hidden layer activation function (i.e., leaky ReLU). The following Equation 1 defines the activations *x*^k^ of the *k*^th^ layer from the activations *x*^k-1^ at the (*k-1*)^th^ layer with the initial activation vector x*^a^*, for a node *h* in the *p*^th^ hidden layer:

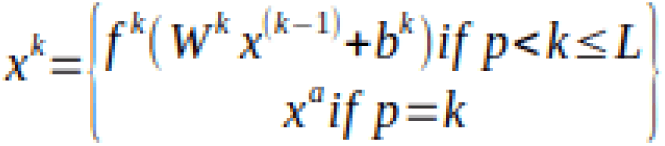

Where f^k^, W^k^, and b^k^ correspond to the k^th^ layer activation function, weight matrix, and bias term, respectively. The activations at the output layer x^L^ have the same dimensions as the model input, and are then used to rank the CpGs in terms of their association with the maximally activated hidden node at layer k-1. The process is then repeated for every hidden node and layer.

### Light-up analysis from inputs

To prioritize CpGs in supervised NCAE-DNN models by their contribution to the output, we applied a perturbation-based forward propagation analysis for feature importance ranking. Since both hyper and hypomethylation are viable states for a CpG site, we first recursively altered input CpG values to either a complete hyper (β = 1) or hypomethylation (β = 0) level, before forward-propagating them through the trained NCAE and DNN. Then, we compared the output of the concatenated models to that of an average methylation profile (mean beta value across samples) representative of a specific tissue and condition, e.g., CD4+ T cells from SLE patients. After the signal propagation was iterated through every input feature (CpG), their contribution to the regression or classification objective of the DNN can be measured by the observed changes in the model outcome, i.e., estimated age or predicted disease classification probability.

### Age estimation using DNAm clocks

We used the getAgeR() function from the R package cgageR to obtain age estimates from Horvath and Hannum DNAm clocks for samples in the multi-tissue (n = 24,676) and the whole blood (n = 13,647) sets. Evaluation metrics were calculated using the functions provided by the R package Metrics. Biologically meaningful age bins were established based on the classification of age categories by the Medical Subject Headings (MeSH) controlled vocabulary thesaurus of the U.S. National Library of Medicine ([http://www.ncbi.nlm.nih.gov/mesh]).

### Gene annotation and enrichment analyses

Genome-wide annotation of CpG probes was performed using Infinium HumanMethylation450 BeadChip probe annotation files and the R package org.Hs.eg.db (version 3.15.0). Pathway and gene ontology enrichment analyses were performed using the respective functions from the R package clusterProfiler (version 4.4.4). Enrichments in disease-associated genes of the top light-up CpG-associated genes per tissue were calculated using a Kolmogorov-Smirnov-like statistic, similar to the enrichment score in gene set enrichment analysis (GSEA) [61]. First, we computed one-sided Fisher’s exact tests for overlap with disease-associated genes retrieved from DisGeNET v7.0 over the cumulative ranked gene lists until the position 1,000. To increase the stringency of the outcome, we avoided the overestimation of the maximal enrichment scores (minimum P) of the light-up ranked gene lists by limiting the minimum cumulative set size to 101 genes. Then, we generated a null distribution of P-values by performing enrichment testing in the same interval across gene lists obtained by randomly assigning gene labels from all CpG-associated genes included in the model input. The procedure was repeated for 1e4 permutations. Finally, we calculated the permutation test P-value by dividing the number of cases in which the null distribution P-values are more extreme than the light-up ranked gene list P-value between the number of permutations. Average permutation P-values were obtained using the function hmp.stat() of the R package harmonicmeanp to compute the harmonic mean P-value for dependent tests.

## DATA AVAILABILITY

The normalized DNA methylation data is available for download at the EWAS Data Hub repository (https://ngdc.cncb.ac.cn/ewas/datahub/repository). The trained models, including the network-coherent AE and supervised deep neural networks NCAE-Age, NCAE-Smoke, and NCAE-SLE are available at https://figshare.com/projects/071350_network_coherent_autoencoders/155090.

## CODE AVAILABILITY

The code used in this study is available at the GitLab repository https://gitlab.com/Gustafsson-lab/deep_methylation_ncaes.

## ACKNOWLEDGMENTS

This work is supported by the Swedish Research Council (grant 2019-04193) and the Wallenberg AI, Autonomous Systems and Software Program (WASP) and SciLifeLab and Wallenberg National Program for Data-Driven Life Science (DDLS) (WASPDDLS21-040/KAW 2020.0239). Computational resources were granted by the Swedish National Infrastructure for Computing (SNIC), including the National Supercomputer Center (NSC, Berzelius-2021-26) and the High Performance Computing Center North (HPC2N, SNIC 2021/5-131 and SNIC 2021/22-199).

## AUTHOR CONTRIBUTIONS

M.G., S.K.D. and D.M. conceived the study and planned the analyses. D.M. performed data pre-processing, model training and evaluation, which were supervised by M.G. with inputs from S.K.D. The manuscript was written by D.M. and M.G. with feedback and inputs from R.J. and S.K.D.

## COMPETING INTERESTS

The authors declare no competing interests.

## Supplementary Data

Supp. File 1: Summary of AE model architecture training and performance.

Supp. File 2: Co-localization and centrality analyses of deep AE latent representations

Suppl. File 3: Co-localization and centrality analyses of sparse AE and variational AE latent representations.

Suppl. File 4: Metadata of EWAS DNA methylation profiles used for AE and supervised model training.

Suppl. File 5: Summary of chronological and biological DNAm age clocks.

Suppl. File 6: Top ranked CpG-associated genes for aging, smoking, and SLE identified using the NCAE-DNN models, by tissue or cell type.

Suppl. File 7: Overlap between aging DNAm signature from the NCAE-Age model and reported DNAm clocks.

Suppl. File 8: Reported DNAm signatures for smoking status.

Suppl. File 9: Pre-processed model input CpG site metadata.

## Notes

https://ngdc.cncb.ac.cn/ewas/datahub/repository

https://figshare.com/projects/071350_network_coherent_autoencoders/155090

https://gitlab.com/Gustafsson-lab/deep_methylation_ncaes

### Competing Interest Statement

The authors have declared no competing interest.

